# Atypical Protein Kinase C iota (PKCλ/ι) Ensures Mammalian Development by Establishing the Maternal-Fetal Exchange Interface

**DOI:** 10.1101/843375

**Authors:** Bhaswati Bhattacharya, Pratik Home, Avishek Ganguly, Soma Ray, Ananya Ghosh, Rashedul Islam, Valerie French, Courtney Marsh, Sumedha Gunewardena, Hiroaki Okae, Takahiro Arima, Soumen Paul

**Author notes:** CORRESPONDING AUTHOR Soumen Paul, University of Kansas Medical Center, 3098 HLSIC, MS 3050, Kansas City, KS 66160, USA.

## Abstract

In utero mammalian development relies on the establishment of the maternal–fetal exchange interface, which ensures transportation of nutrients and gases between the mother and the fetus. This exchange interface is established via development of multinucleated syncytiotrophoblast cells (SynTs) during placentation. In mouse, SynTs develop via differentiation of the trophoblast progenitor cells (TSPCs) of the placenta primordium and in human, SynTs are developed via differentiation of villous cytotrophoblast (CTB) progenitors. Despite the critical need in pregnancy progression, conserved signaling mechanisms that ensure SynT development are poorly understood. Herein, we show that Atypical Protein Kinase C iota (PKCλ/I) plays an essential role in establishing the SynT differentiation program in trophoblast progenitors. Loss of PKCλ/I in the mouse TSPCs abrogates SynT development leading to embryonic death at ~E9.0. We also show that PKCλ/I-mediated priming of trophoblast progenitors for SynT differentiation is a conserved event during human placentation. PKCλ/I is selectively expressed in the first-trimester CTBs of a developing human placenta. Furthermore, loss of PKCλ/I in CTB-derived human trophoblast stem cells (Human TSCs) impairs their SynT differentiation potential both *in vitro* and after transplantation in immunocompromised mice. Our mechanistic analyses indicate that PKCλ/I signaling maintains expression of GCM1, GATA2, and PPARγ, which are key transcription factors to instigate SynT differentiation programs in both mouse and human trophoblast progenitors. Our study uncovers a conserved molecular mechanism, in which PKCλ/I signaling regulates establishment of the maternal-fetal exchange surface by promoting trophoblast progenitor to SynT transition during placentation.

## Introduction

Trophoblast progenitors are critical for embryo implantation and early placentation. Defective development and differentiation of trophoblast progenitors during early human pregnancy either leads to pregnancy failure (Cockburn & Rossant, 2010; Pfeffer & Pearton, 2012; R. M. Roberts & S. J. Fisher, 2011; J. Rossant & J. C. Cross, 2001), or pregnancy-associated complications like fetal growth retardation and preeclampsia (Myatt, 2006; Pfeffer & Pearton, 2012; Redman & Sargent, 2005; J. Rossant & J. C. Cross, 2001), or serves as the developmental cause for postnatal or adult diseases (Funai et al., 2005; Gluckman, Hanson, Cooper, & Thornburg, 2008; Godfrey & Barker, 2000). However, due to experimental and ethical barrier, we have a poor understanding of molecular mechanisms that are associated with early stages of human placentation. Rather, gene knockout studies in mice have provided important information about molecular mechanisms that regulate mammalian placentation. While mouse and human placentae differ in their morphology and trophoblast cell types, important similarities exist in the formation of the maternal-fetal exchange interface. Both mice and humans display hemochorial placentation (Carter, 2007), where the maternal-fetal exchange interface is established via direct contact between maternal blood and placental SynTs.

In a peri-implantation mouse embryo, proliferation and differentiation of polar trophectoderm results in the formation of TSPCs, which reside within the extra-embryonic ectoderm ExE (Rossant, 2001), and later in ExE-derived ectoplacental cone (EPC) and chorion. Subsequently, the TSPCs within the ExE/EPC region contribute to develop the junctional zone, a compact layer of cells sandwiched between the labyrinth and the outer TGC layer. Development of the junctional zone is associated with differentiation of trophoblast progenitors to four trophoblast cell lineages (i) TGCs (Simmons, Fortier, & Cross, 2007), (ii) spongiotrophoblast cells (Simmons & Cross, 2005), (iii) glycogen cells and (iv) invasive trophoblast cells that invade the uterine wall and maternal vessels (Kaufmann, Black, & Huppertz, 2003; Rosario, Konno, & Soares, 2008; Soares et al., 2012).

The mouse placental labyrinth, which constitutes the maternal-fetal exchange interface, develops after the allantois attaches with the chorion. The multilayered chorion forms around embryonic day (E) 8.0 when chorionic ectoderm fuses to basal EPC, thereby reuniting TSPC populations separated by formation of the ectoplacental cavity (Simmons et al., 2008). Subsequently, the chorion attaches with the allantois to initiate the development of the placental labyrinth, which contains two layers of SynTs, known as SynT-I and SynT-II. At the onset of labyrinth formation, Glial Cells Missing 1 (*Gcm1*) expression is induced in the TSPCs of the chorionic ectoderm (Basyuk et al., 1999), which promotes cell cycle exit and differentiation to the SynT-II lineage (Simmons et al., 2008). Whereas the TSPCs of the basal EPC progenitors that express Distal-less 3 (*Dlx3*), contributes to syncytial SynT-I lineage (Simmons et al., 2008).

In contrast to mouse, the earliest stage of human placentation is associated with the formation of a zone of invasive primitive syncytium at the blastocyst implantation site (Boss, Chamley, & James, 2018; James, Carter, & Chamley, 2012; Knofler et al., 2019). Later, columns of CTB progenitors penetrate the primitive syncytium to form primary villi. With the progression of pregnancy, primary villi eventually branch and mature to form the villous placenta, containing two types of matured villi; (i) anchoring villi, which anchor to maternal tissue and (ii) floating villi, which float in the maternal blood of the intervillous space (Boss et al., 2018; James et al., 2012; Knofler et al., 2019). The proliferating CTBs within anchoring and floating villi adapt distinct differentiation fates during placentation (Haider et al., 2016). In anchoring villi, CTBs establish a column of proliferating CTB progenitors known as column CTBs (Haider et al., 2016), which differentiate to invasive extravillous trophoblasts (EVTs), whereas, CTB progenitors of floating villi (villous CTBs) differentiate and fuse to form the outer multinucleated SynT layer. The villous CTB-derived SynTs establish the nutrient, gas and waste exchange surface, produce hormones and promote immune tolerance to fetus throughout gestation (Beer & Sio, 1982; Chamley et al., 2014; Cole, 2012; Costa, 2016; PrabhuDas et al., 2015; Yang, Lei, & Rao Ch, 2003).

Thus, the establishment of the placental exchange surface in both mouse and human are associated with the formation of differentiated, multinucleated SynTs from the trophoblast progenitors of placenta primordia. Moreover, both mouse and human trophoblast progenitors express key transcription factors like, GCM1, DLX3, Peroxisome proliferator-activated nuclear receptor gamma (PPARγ) and GATA binding protein 2 (GATA2), which are shown to be important for SynT development during placentation (Anson-Cartwright et al., 2000; Cockburn & Rossant, 2010; Pratik Home et al., 2017; Parast et al., 2009; Pfeffer & Pearton, 2012; R. M. Roberts & S. J. Fisher, 2011; Rossant, 2004; Janet Rossant & James C. Cross, 2001; Stecca et al., 2002). Despite these similarities, conserved signaling pathways that program SynT development in both mouse and human trophoblast progenitors are incompletely understood. Fortunately, the success in deriving true human TSCs from villous cytotrophoblast cells (CTBs) (Okae et al., 2018) have opened up new possibilities for direct assessment of conserved mechanisms that prime differentiation of multipotent trophoblast progenitor to SynT lineage. Therefore, we herein analyzed both mouse mutants and human TSCs to test the specific role of PKCλ/I in that process.

The PKCλ/I belongs to the atypical group of PKCs, which consists of another isoform PKCζ. The aPKC isoforms have been implicated in cell lineage patterning in preimplantation embryos (Zhu, Leung, Shahbazi, & Zernicka-Goetz, 2017). We demonstrated that both PKCζ and PKCλ/I regulate self-renewal vs. differentiation potential in both mouse and rat embryonic stem cells (Dutta et al., 2011; Mahato et al., 2014; Rajendran et al., 2013). Interestingly, gene knockout studies in mice indicated that PKCζ is dispensable for embryonic development (Leitges et al., 2001), whereas ablation of PKCλ/I results in early gestation abnormalities leading to embryonic lethality (Seidl et al., 2013; Soloff, Katayama, Lin, Feramisco, & Hedrick, 2004) prior to embryonic day (E) 9.5, a developmental stage equivalent to first trimester in humans. However, importance of PKCλ/I in the context of post-implantation trophoblast lineage development during mouse or human placentation has never been addressed. We found that PKCλ/I protein is specifically abundant in TSPCs and villous CTBs within the developing mouse and human placenta, respectively. We show that both global and trophoblast-specific loss of PKCλ/I in a mouse embryo is associated with defective development of placental labyrinth due to impairment of gene expression programming that ensures SynT development. We further demonstrate that the PKCλ/I-signaling in human TSCs is also essential for maintaining their SynT differentiation potential. Our analyses revealed an evolutionarily conserved, developmental stage-specific mechanism in which PKCλ/I-signaling orchestrates gene expression program in trophoblast progenitors for successful progression of in utero mammalian development.

## RESULTS

### PKCλ/I protein expression is selectively abundant in the trophoblast progenitors of developing mouse and human placentae

Earlier studies showed that PKCλ/I is ubiquitously expressed in all cells of a developing pre-implantation mouse embryo (Saiz, Grabarek, Sabherwal, Papalopulu, & Plusa, 2013), including the trophectoderm cells. Also, another study showed that PKCλ/I is ubiquitously expressed in both embryonic and extraembryonic cell lineages in a postimplantation E7.5 mouse embryo (Seidl et al., 2013). We validated both of these observations in a blastocyst (Fig. S1A) and in E7.5 mouse embryo (Fig. S1B). However, relative abundance of PKCλ/I protein expression in different cell types is not well documented during postimplantation mouse development. Therefore, we tested PKCλ/I protein expression at different stages of mouse post-implantation development. We found that in ~E8 mouse embryos, PKCλ/I protein is most abundantly expressed in the TSPCs residing in the placenta primordium (Fig. 1A). In comparison, cells within the developing embryo proper showed very low levels of PKCλ/I protein expression. The high abundance of *Prkci* mRNA and PKCλ/I protein expression was detected in the trophoblast cells of an E9.5 mouse embryo (Fig. S1C and Fig. 1B). We also observed that PKCλ/I is expressed in the maternal decidua. Surprisingly, the embryo proper showed extremely low PKCλ/I protein expression at this developmental stage. Furthermore, the selective abundance of PKCλ/I protein expression in the trophoblast is also maintained as development progresses to mid-gestation as in E12.5 (Fig. S1D).

**Figure 1:**
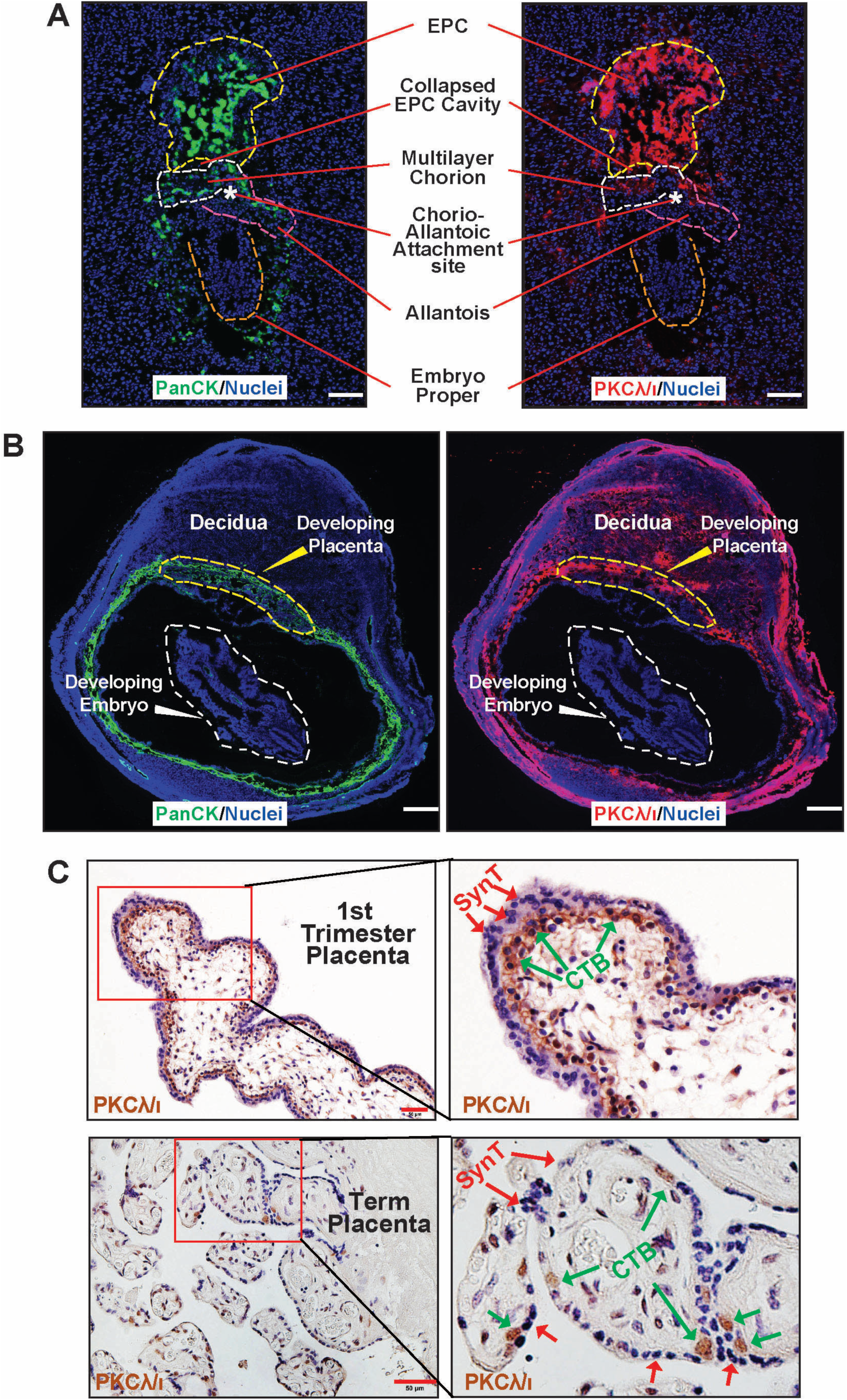
PKCλ/I protein expression is selectively abundant in trophoblast progenitors of early postimplantation mammalian embryos. (A) Immunofluorescence images showing trophoblast progenitors, marked by anti-pancytokeratin antibody (green), expressing high levels of PKCλ/I protein (red) in an ~E8 mouse embryo. Note much less expression of PKCλ/I protein within cells of the developing embryo proper (orange dotted boundary). EPC, Chorion, allantois and the chorio-allantoic attachment site are indicated with yellow, white, pink dotted lines and a white asterisk, respectively. (B) Immunofluorescence images of an E9.5 mouse implantation sites showing pan-Cytokeratin (left), PKCλ/I (right) and nuclei (DAPI). At this developmental stage, PKCλ/I protein is highly expressed in trophoblast cells of the developing placenta and in maternal uterine cells. However, PKCλ/I protein expression is much less in the developing embryo (C) Immunohistochemistry showing PKCλ/I is selectively expressed within the cytotrophoblast progenitors (green arrows) of first-trimester (8 week) and term (38 week) human placentae.

As PKCλ/I expression during human placentation has never been tested, we investigated PKCλ/I protein expression in human placentae at different stages of gestation. Our analyses revealed that PKCλ/I protein is expressed specifically in the villous CTBs within a first-trimester human placenta (Fig. 1C). PKCλ/I is also expressed at a reduced level in CTBs within term placenta (Fig. 1C). Intriguingly, differentiated SynTs in first-trimester human placentae as well as in term-placentae lack PKCλ/I protein expression. Thus, our analyses revealed a selective abundance of PKCλ/I protein expression in trophoblast progenitors of both mouse and human placentae.

### Global Loss of PKCλ/I in a developing mouse embryo abrogates placentation

Global deletion of Prkci in a developing mouse embryo (PKCλ/I KO embryo) results in gastrulation defect leading to embryonic lethality (Seidl et al., 2013; Soloff et al., 2004) at ~E9.5. However, the placentation process was never studied in PKCλ/I KO mouse embryos. As many embryonic lethal mouse mutants are associated with placentation defects, we probed into trophoblast development and placentation in post-implantation PKCλ/I KO embryos. We started investigating placenta and trophoblast development in PKCλ/I KO embryos starting from E7.5. At this stage the placenta primordium consists of the ExE/EPC regions. However, we did not notice any obvious phenotypic differences of the ExE/EPC development between the control and the PKCλ/I KO embryos at E7.5 (Fig. 2A, left panel and Fig. S2A). We noticed developmental defect in PKCλ/I KO placentae after the chorio-allantoic attachment, an event which takes place at ~ E8.5. We noticed that PKCλ/I KO developing placentae were smaller in size at E8.5 (Fig. 2A, middle panel) and this defect in placentation was more prominent in E9.5 embryos. At E9.5, the placentae in PKCλ/I KO embryos were significantly smaller in size (Fig. 2A, right panel and Fig. S2A) and the embryo proper showed gross impairment in development due to defective gastrulation (Fig. 2B), as reported in earlier studies (Soloff et al., 2004). Immunofluorescence analyses revealed defective embryonic-extraembryonic attachment with an altered orientation of the embryo proper with respect to the developing placenta (Fig. 2C). We also failed to detect any visible labyrinth zone in PKCλ/I KO placentae. The abrogation of the placental labyrinth and the SynT development were confirmed by a near complete absence of *Gcm1* mRNA expression (Fig. S2B) and lack of any Dlx3-expressing labyrinth trophoblast cells in E9.5 PKCλ/I KO placentae (Fig. 2D). However, the trophoblast giant cells (TGCs) developed and were present in E9.5 PKCλ/I KO placentae. We could not test placentation in the PKCλ/I KO embryos beyond E9.5 as these embryos and placentae begin to resorb at late gestational stages. Thus, from our findings, we concluded that the global loss of PKCλ/I in a developing mouse embryo leads to defective placentation after the chorio-allantoic attachment due to impaired development of the SynT lineage, resulting in abrogation of placental labyrinth formation.

**Figure 2:**
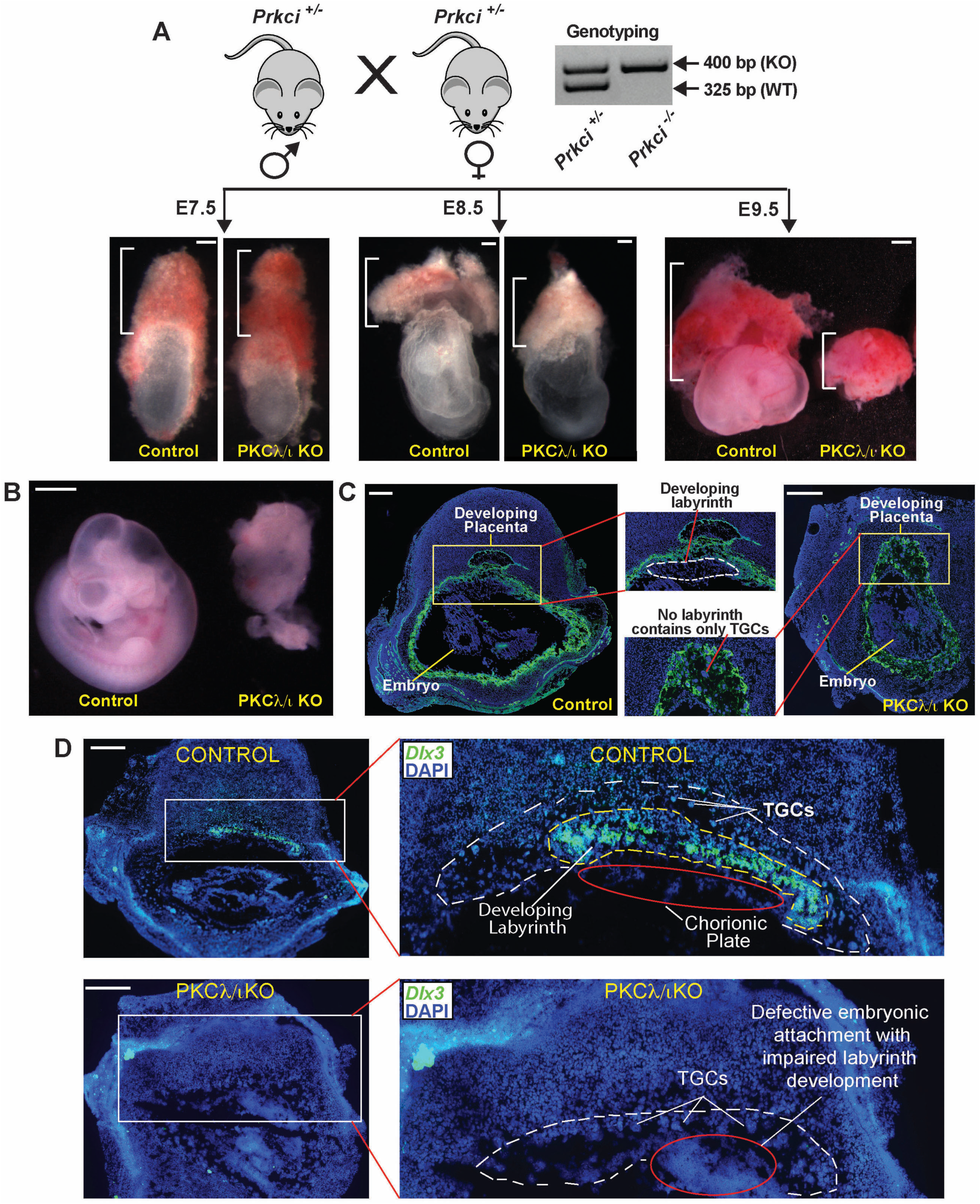
Global Loss of PKCλ/I in a developing mouse embryo abrogates placentation. (A) Experimental strategy and phenotype of mouse conceptuses defining the importance of PKCλ/I in placentation. Heterozygous (*Prkci*^+*/*−^) male and female mice were crossed to generate homozygous knockout (*Prkci*^−*/*−^, PKCλ/I KO) embryos and confirmed by genotyping. Embryonic and placental developments were analyzed at E7.5, E8.5 and E9.5 and representative images are shown. At E7.5, placenta primordium developed normally in PKCλ/I KO embryos. However, defect in placentation in PKCλ/I KO conceptuses were observable (smaller placentae) at E8.5 and was prominent at E9.5. (B) Developing control and PKCλ/I KO embryos were isolated at ~E9.5 and representative images are shown. The PKCλ/I KO embryo proper shows gastrulation defect as described in earlier study. (C) Placentation at control and PKCλ/I KO implantation sites were analyzed at ~E9.5 via immunostaining with anti-pan-cytokeratin antibody (green, trophoblast marker). The developing PKCλ/I KO placenta lacks the labyrinth zone and mainly contains the TGCs (red line). Also, unlike in control embryos, the developmentally arrested PKCλ/I KO placenta and embryo proper are not segregated and are attached together. (D) RNA in situ hybridization assay was performed using fluorescent probes against *Dlx3* mRNA. Images show that, unlike the control placenta, the PKCλ/I KO placenta lacks *Dlx3* expressing labyrinth trophoblast cells.

### Trophoblast-specific PKCλ/I depletion impairs mouse placentation leading to embryonic death

Since we observed placentation defect in the global PKCλ/I KO embryos, we next interrogated the importance of trophoblast cell-specific PKCλ/I function in mouse placentation and embryonic development. Although a *Prkci*-conditional knockout mouse model exists, we could not get access to that mouse. Therefore, we performed RNAi using lentiviral mediated gene delivery approach as described earlier (Lee, Rumi, Konno, & Soares, 2009) to specifically deplete PKCλ/I in the developing trophoblast cell lineage. We transduced zona-removed mouse blastocysts with lentiviral particles with shRNA against *Prkci* (Fig. 3A) and transferred them to pseudo pregnant females. We confirmed the efficiency of shRNA-mediated PKCλ/I-depletion by measuring *Prkci* mRNA expression in transduced blastocysts (Fig. 3B) and also by testing loss of PKCλ/I protein expressions in trophoblast cells of developing placentae (Fig. S3). Intriguingly, the trophoblast-specific PKCλ/I depletion also resulted in embryonic death before E9.5 due to severe defect in embryo patterning (Fig. 3C and 3D). Furthermore, the immunofluorescence analyses of trophoblast cells at ~E9.5 confirmed defective placentation in the *Prkci* KD placentae, characterized with a near complete absence of the labyrinth zone (Fig. 3E) and *Dlx3* expressing SynT populations (Fig. 3F). Thus, the trophoblast-specific depletion of PKCλ/I in a developing mouse embryo recapitulated similar placentation defect and embryonic death as observed in the global PKCλ/I KO embryos.

**Figure 3:**
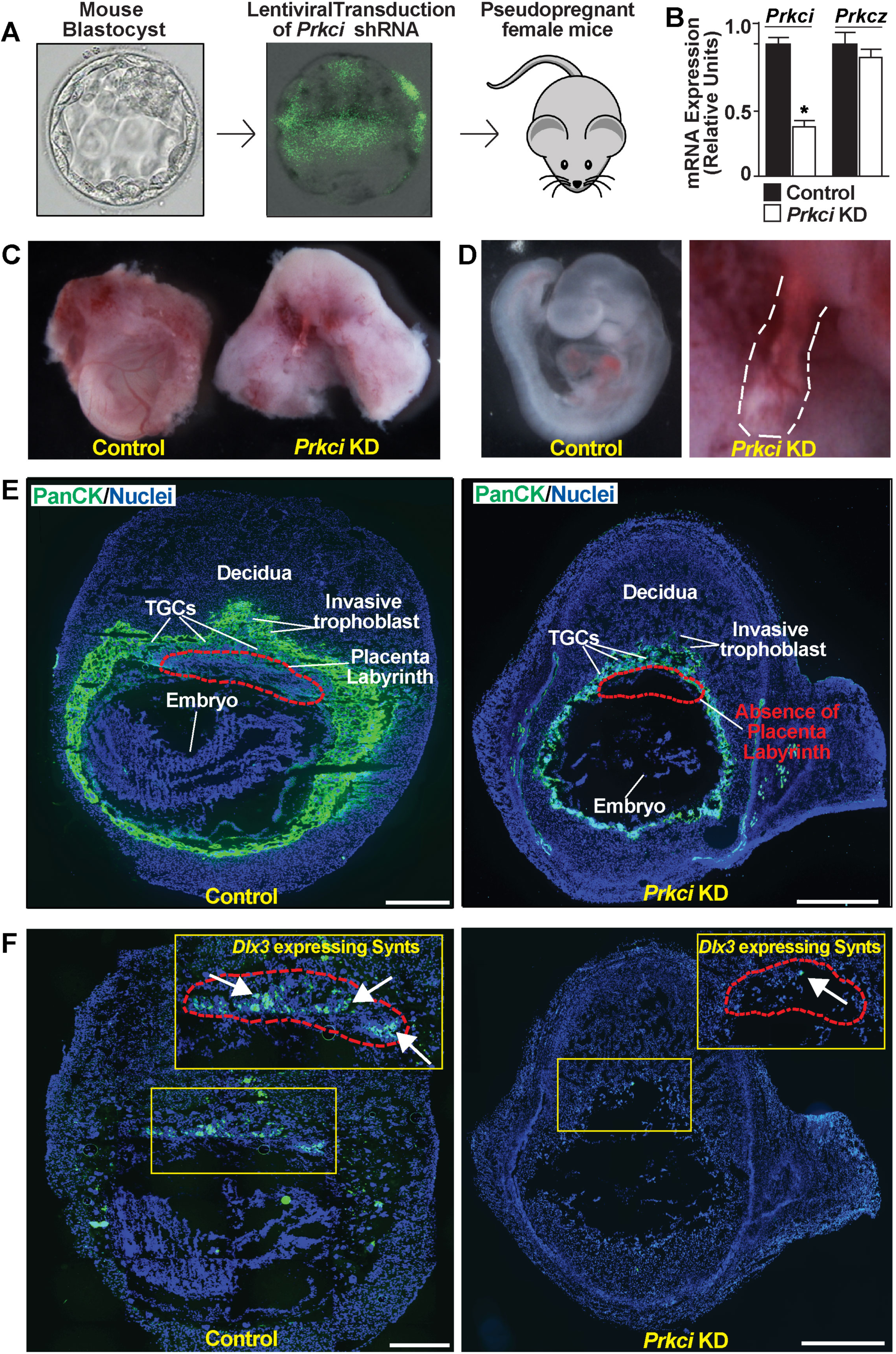
Trophoblast-specific PKCλ/I depletion impairs mouse placentation leading to embryonic death. (A) Schematics to study developing mouse embryos with trophoblast-specific depletion of *Prkci* (*Prkci* KD embryos). Blastocysts were transduced with lentiviral vectors expressing shRNA against *Prkci* and transduction was confirmed by monitoring EGFP expression. Transduced blastocysts were transferred into the uterine horns of pseudo pregnant females to study subsequent effect on embryonic and placental development. (B) Knockdown efficiency with shRNA was confirmed by testing loss of *Prkci* mRNA expression in transduced blastocysts. (C) and (D) Representative images show control and *Prkci* KD conceptuses and developing embryo proper, isolated at E9.5. Similar to global PKCλ/I KO embryos, trophoblast-specific *Prkci* KD embryos showed severe developmental defect. (E) Immunostaining with anti-pan-cytokeratin antibody (green, trophoblast marker) showed defective placentation in the *Prkci* KD implantation sites at ~E9.5. The images show that unlike the control placenta, labyrinth formation was abrogated in the *Prkci* KD placenta. (F) RNA in situ hybridization assay confirmed near-complete absence of *Dlx3* expressing trophoblast cells in the *Prkci* KD placenta.

### PKCλ/I signaling in a developing mouse embryo is essential to establish a transcriptional program for TSPC to SynT differentiation

The abrogation of labyrinth development in the trophoblast-specific *Prkci* KD mouse placentae indicated a critical importance of the PKCλ/I signaling in SynT development and labyrinth formation. During mouse placentation, the SynT differentiation is associated with the suppression of TSC/TSPC-specific genes, such as Caudal type homeobox 2 (*Cdx2)*, Eomesodermin *(Eomes)*, TEA domain transcription factor 4 *(Tead4)*, Estrogen related receptor beta *(Esrrβ)* and E74 like transcription factor 5 (*Elf5)* (Donnison et al., 2005; P. Home et al., 2012; Latos et al., 2015; R. Michael Roberts & Susan J. Fisher, 2011; Russ et al., 2000; Strumpf et al., 2005) and induction of expression of the SynT-specific genes, such as *Gcm1, Dlx3* and fusogenic retroviral genes SyncytinA and SyncytinB (Janet Rossant & James C. Cross, 2001). In addition, other transcription factors, such as PPARγ, GATA transcription factors-GATA2 and GATA3, and cell signaling regulators, including members of mitogen activated protein kinase pathway are implicated in mouse SynT development (Pratik Home et al., 2017; Nadeau et al., 2009; Parast et al., 2009). Therefore, to define the molecular mechanisms of PKCλ/I-mediated regulation of SynT development, we specifically depleted PKCλ/I expression in mouse TSCs via RNAi (Fig. 4A) and asked whether the loss of PKCλ/I impairs mouse TSC self-renewal or their differentiation to specialized trophoblast cell types.

**Figure 4:**
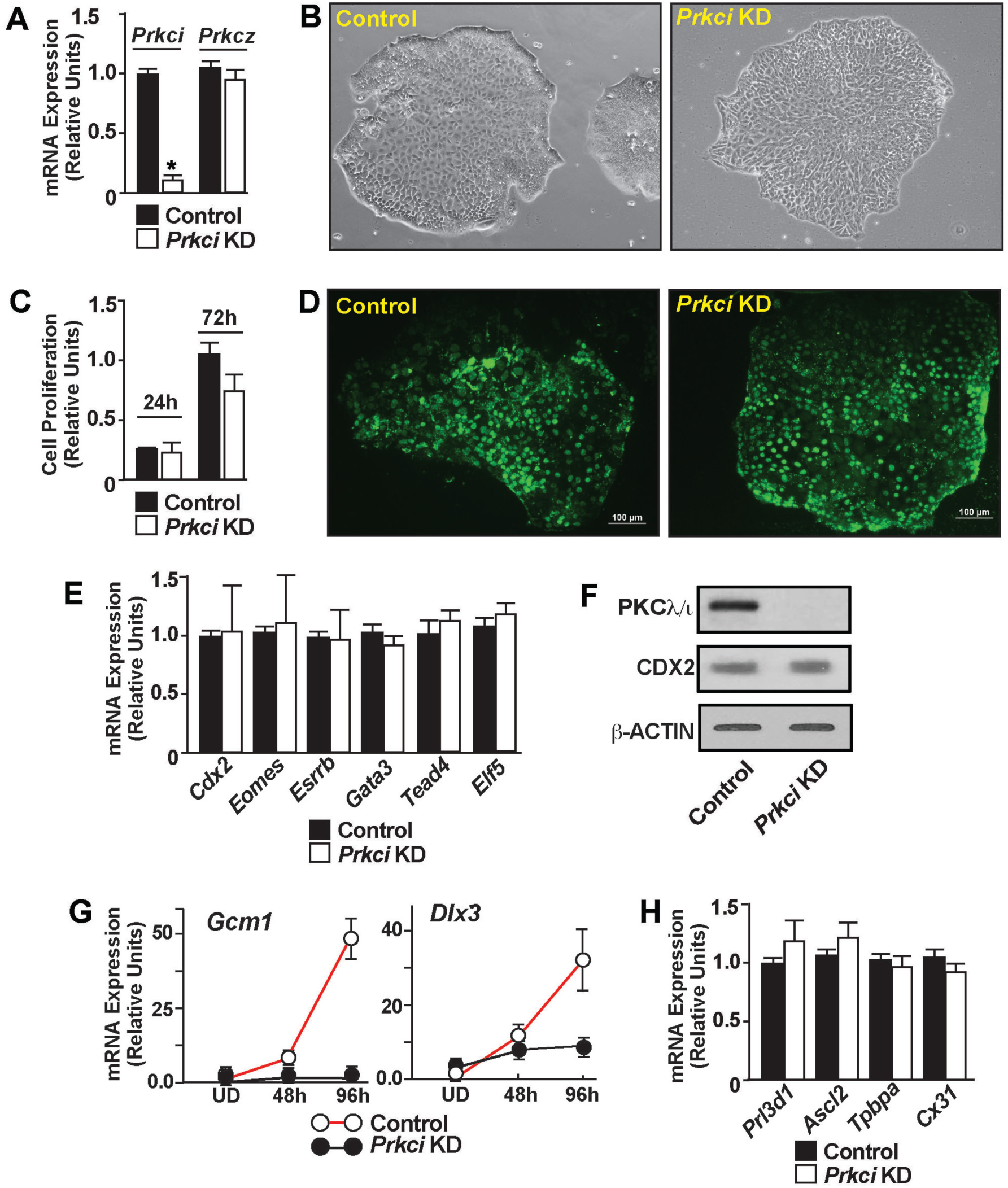
PKCλ/I signaling is essential to establish a transcriptional program for SynT differentiation. (A) Quantitative RT-PCR to validate loss of *Prkci* mRNA expression in *Prkci* KD mouse TSCs (mean ± SE; n = 3, p≤0.001). The shRNA molecules targeting the *Prkci* mRNA had no effect on *Prkcz* mRNA expression (B) Morphology of control and *Prkci* KD mouse TSCs. (C) and (D) Assessing proliferation rate of control and *Prkci* KD mouse TSCs by MTT assay and BrdU labeling, respectively. (E) Quantitative RT-PCR (mean ± SE; n = 3), showing unaltered mRNA expression of trophoblast stem-state specific genes like *Cdx2, Eomes, Tead4, Gata3, Elf5 and Esrrb* in *Prkci* KD mouse TSCs. (F) Western blot analyses confirming unaltered CDX2 protein expression in mouse TSCs upon PKCλ/I depletion. (G) *Prkci* KD mouse TSCs were allowed to grow under differentiation conditions and gene expression analyses of syncytiotrophoblast markers *Gcm1* and *Dlx3* were done (mean ± SE; n = 3). The plots show that loss of PKCλ/I in mouse TSCs results in impaired induction of *Gcm1* and *Dlx3* expression. (H) Quantitative RT-PCR analyses of *Prl3d1, Ascl2, Tpbpa* and *Cx31* mRNA expression in differentiated *Prkci* KD mouse TSCs. (mean ± SE; n = 3) to assess TGCs, spongiotrophoblasts and glycogen trophoblast cell differentiation, respectively. Expression of these genes were not altered in differentiated *Prkci* KD mouse TSCs, indicating PKCλ/I is dispensable for mouse TSC differentiation to TGCs, spongiotrophoblasts and glycogen trophoblast cells.

When cultured in stem-state culture condition (with FGF4 and Heparin), PKCλ/I-depleted mouse TSCs (*Prkci* KD mouse TSCs) did not show any defect in the stem-state colony morphology (Fig. 4B). Also, cell proliferation analyses by MTT assay and BrDU incorporation assay indicated that cell proliferation was not affected in the *Prkci* KD mouse TSCs (Fig. 4C and 4D). Furthermore, mRNA expression analyses showed that expression of TSC stem-state regulators, such as *Cdx2, Eomes, Gata3, Tead4, Esrrb and Elf5* were not affected upon PKCλ/I depletion (Fig. 4E). Western blot analysis also confirmed that CDX2 protein expression was not affected in the *Prkci* KD TSCs (Fig. 4F). These results indicated that PKCλ/I signaling is not essential to maintain the self-renewal program in mouse TSCs.

Next, we asked whether the loss of PKCλ/I affects mouse TSC differentiation program. Removal of FGF4 and heparin from the culture medium induces spontaneous differentiation in mouse TSCs, which can be monitored over a course 6-8 days. During this differentiation program, induction of SynT-differentiation markers like *Gcm1, Dlx3* can be monitored in differentiating cells between day 2 - day 4. Subsequently, the TSC markers are repressed in the differentiating TSCs as the TGC-specific differentiation program becomes more prominent. Thus, after day 6 of differentiation, mouse TSCs highly express TGC specific markers, like Prolactin family 3 subfamily d member 1 (*Prl3d1)*, Heart and neural crest-derived transcript 1 *(Hand1)*, Prolactin family 2 subfamily c member 2 *(Prl2c2)*. In addition, Trophoblast specific protein alpha (*Tpbpa)*, Achaete-scute homologue 2 *(Ascl2)*, Connexin 31 *(Cx31)*, which are markers of spongiotrophoblast and glycogen trophoblast cells of the placental junctional zone, are also induced in differentiated mouse TSCs.

As the loss of PKCλ/I affects labyrinth development, we monitored expressions of *Gcm1 and Dlx3* in differentiating *Prkci* KD mouse TSCs. Similar to our findings with the PKCλ/I-depleted placentae, induction of *Dlx3* and *Gcm1* mRNA expression was impaired in differentiating *Prkci* KD mouse TSCs (Fig. 4G). In contrast, induction of *Tpbpa, Prl3d1, Cx31*, which are markers for spongiotrophoblasts, TGCs and glycogen cells, respectively, were not affected in differentiated *Prkci* KD mouse TSCs (Fig. 4H). Thus, we concluded that the loss of PKCλ/I in mouse TSCs does not affect their differentiation to specialized trophoblast cells of the placental junctional zone, such as spongiotrophoblasts, TGCs, and the glycogen cells. Rather, PKCλ/I is essential to specifically establish the SynT differentiation program in mouse TSCs.

Based on our findings with *Prkci* KD mouse TSCs, we hypothesized that the PKCλ/I signaling might regulate key genes, which are specifically required to induce the SynT differentiation program in TSCs. To test this hypothesis, we performed unbiased whole RNA sequencing (RNA-seq) analysis with *Prkci* KD mouse TSCs. RNA-seq analyses showed that the depletion of PKCλ/I in mouse TSCs altered expression of 164 genes by at least two folds with a high significance level (p≤0.01). Among these 164 genes, 120 genes were down-regulated and 44 genes were up-regulated (Fig. 5A and Supplemental Tables S1 and S2). Ingenuity pathway analyses revealed multi-modal biofunctions of PKCλ/I regulated genes, including involvement in embryonic and reproductive developments (Fig. 5B). To further gain confidence on PKCλ/I regulated genes in the mouse TSCs, we curated the number of altered genes with a false discovery rate (FDR) threshold of 0.1. The FDR filtering identified only 6 upregulated genes and 46 downregulated genes in the *Prkci* KD mouse TSCs (Fig. 5C, 5D and Supplemental Table S3). Among the downreguated genes, *Prkci* was identified as the most significantly altered gene, thereby confirming the specificity and high efficiency of the shRNA-mediated *Prkci* depletion.

**Figure 5:**
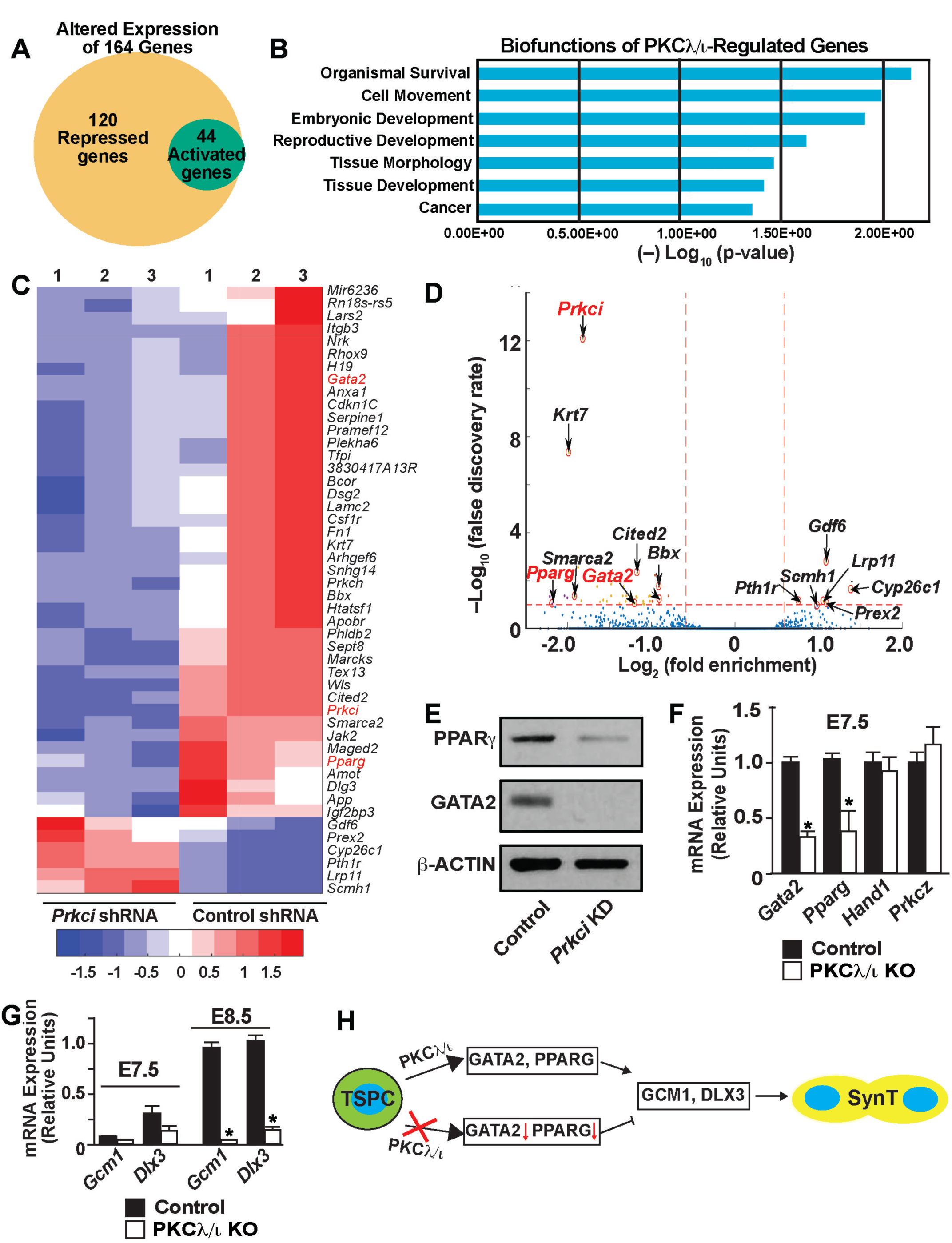
PKCλ/I signaling regulates expression of GATA2 and PPARγ, key transcription factors for SynT differentiation, in mouse TSCs and primary TSPCs of a mouse placenta primordium. (A) Whole genome RNA-seq was performed in *Prkci* KD mouse TSCs and the Venn diagram shows number of genes that were significantly downregulated (120 genes) and upregulated (44 genes) upon PKCλ/I depletion. (B) The plot shows most significant biofunctions (identified via Ingenuity Pathway Analysis) of PKCλ/I-regulated genes in mouse TSCs. (C) and (D) Heat map and Volcano plot, respectively, showing significantly altered genes in *Prkci* KD mouse TSCs. Along with *Prkci, Gata2* and *Pparg* (marked in red) are among the most significantly down-regulated genes in *Prkci* KD mouse TSCs. (E) Western blot analyses showing loss of GATA2 and PPARγ protein expressions in *Prkci* KD mouse TSCs. (F) Quantitative RT-PCR analyses showing downregulation of *Gata2* and *Pparg* mRNA expression in TSPCs of E7.5 PKCλ/I KO placenta primordium (mean ± SE; n = 4, p≤0.01). Expression of *Hand1* and *Prkcz* mRNAs remain unaltered. (G) Quantitative RT-PCR analyses in E7.5 and E8.5 PKCλ/I KO placenta primordia (mean ± SE; n = 4, p≤0.001) showing abrogation of *Gcm1* and *Dlx3* induction, which happens between E7.5 and E8.5, in PKCλ/I KO developing placentae. (H) The model implicates a PKCλ/I-GATA2/PPARγ-GCM1/DLX3 regulatory axis in SynT differentiation during mouse placentation.

Among the six genes, which were significantly upregulated in *Prkci* KD mouse TSCs, only Growth differentiation factor 6 *(Gdf6)* has been implicated in trophoblast biology in an overexpression experiment with embryonic stem cells (Lichtner, Knaus, Lehrach, & Adjaye, 2013). However, *Gdf6* deletion in a mouse embryo does not affect placentation (Clendenning & Mortlock, 2012; Mikic, Rossmeier, & Bierwert, 2009). In contrast, three transcription factors, *Gata2, Pparg* and *Cited2*, which were significantly downregulated in the *Prkci* KD mouse TSCs, are implicated in the regulation of trophoblast differentiation and labyrinth development. Earlier gene knockout studies implicated CITED2 in the placental labyrinth formation. However, CITED2 is proposed to have a non-cell autonomous role in SynT as its function is more important in proper patterning of embryonic capillaries in the labyrinth zone rather than in promoting the SynT differentiation (Withington et al., 2006). In contrast, knockout studies in mouse TSCs indicated that PPARγ is an important regulator for SynT differentiation (Parast et al., 2009). PPARγ-null mouse TSCs showed specific defects in SynT differentiation and rescue of PPARγ expression rescued *Gcm1* expression and SynT differentiation. Also, earlier we showed that in mouse TSCs, GATA2 directly regulates expression of several SynT-associated genes including *Gcm1* and, in coordination with GATA3, ensures placental labyrinth development (Pratik Home et al., 2017). Therefore, we focused our study on GATA2 and PPARγ and further tested their expressions in *Prkci* KD mouse TSCs and PKCλ/I KO placenta primordium. We validated the loss of GATA2 and PPARγ protein expressions in *Prkci* KD mouse TSCs (Fig. 5E). Our analyses confirmed that both *Gata2* and *Pparg* mRNA expression are significantly downregulated in E7.5 PKCλ/I KO placenta primordium (Fig. 5F) and the loss of *Gata2* and *Pparg* expression was subsequently associated with impaired transcriptional induction of both *Gcm1 and Dlx3* in E8.5 PKCλ/I KO placentae (Fig. 5G). Thus, our studies in *Prkci* KD mouse TSCs and PKCλ/I KO placenta primordium indicated a regulatory pathway, in which the PKCλ/I signaling in differentiating TSPCs ensures GATA2 and PPARγ expression, which in turn establish proper transcriptional program for SynT differentiation (Fig. 5H).

### PKCλ/I is critical for human trophoblast progenitors to undergo differentiation towards syncytiotrophoblast lineage

Our expression analyses revealed that PKCλ/I expression is conserved in CTB progenitors of a first-trimester human placenta. However, functional importance of PKCλ/I in the context of human trophoblast development and function has never been tested. We wanted to test whether PKCλ/I signaling mediates a conserved function in human CTB progenitors to induce SynT differentiation. However, testing molecular mechanisms in isolated primary first-trimester CTBs is challenging due to lack of established culture conditions, which could maintain CTBs in a self-renewing stage or could promote their differentiation to SynT lineage. Rather, the recent success of derivation of human trophoblast stem cells (human TSCs) from first-trimester CTBs (Okae et al., 2018) has opened up new opportunities to define molecular mechanisms that control human trophoblast lineage development. When grown in media containing a Wnt activator CHIR99021, EGF, Y27632 (a Rho-associated protein kinase [ROCK] inhibitor), A83-01, and SB431542 (TGF-β inhibitors) and valproic acid (VPA) (a histone deacetylase [HDAC] inhibitor), the established human TSCs can be maintained in a self-renewing stem state for multiple passages. In contrast, when cultured in the presence of cAMP agonist forskolin, human TSCs synchronously differentiate and fuse to form two-dimensional (2D) syncytia on a high-attachment culture plate or three-dimensional cyst-like structures on a low adhesion culture plate. In both 2D and 3D culture conditions, differentiated human TSCs highly express SynT markers and secrete a large amount of human chorionic gonadotropin (hCG). Thus, depending on the culture conditions, human TSCs efficiently recapitulate both the self-renewing CTB progenitor state and their differentiation to SynTs. Therefore, we used human TSCs as a model system and performed loss-of-function analyses to test importance of PKCλ/I signaling in human TSC self-renewal vs. their differentiation towards the SynT lineage.

We performed lentiviral-mediated shRNA delivery to deplete PKCλ/I expression in human TSCs (*PRKCI* KD human TSCs). The shRNA-mediated RNAi in human TSCs reduced *PRKCI* mRNA expression by more than 90% without affecting the *PRKCZ* mRNA expression (Fig. 6A and 6B), thereby, confirming the specificity of the RNAi approach. Similar to *Prkci* KD mouse TSCs, *GATA2, PPARG* and *GCM1* mRNA expressions were significantly reduced in *PRKCI* KD human TSCs (Fig. 6A). Western blot analyses also confirmed loss of GATA2 and PPARγ protein expressions in *PRKCI* KD human TSCs (Fig. 6B). However, the loss of PKCλ/I expression in human TSCs did not overtly affect their proliferation (Fig. S4B and S4C), stem-state morphology (Fig. 6C and S4A) or expression of trophoblast stem state markers, such as *TEAD4 or CDX2* (Fig. S4D). Rather, we observed a smaller (~25%) induction in *ELF5* mRNA expression upon loss of PKCλ/I (Fig. S4D). Thus, we concluded that PKCλ/I signaling is not essential to maintain the self-renewing stem-state in human TSCs, rather it is important to maintain optimum expression of key genes like *GATA2, PPARG* and *GCM1*, which are known regulators of SynT differentiation. Therefore, we next interrogated SynT differentiation efficiency in *PRKCI* KD human TSCs.

**Figure 6:**
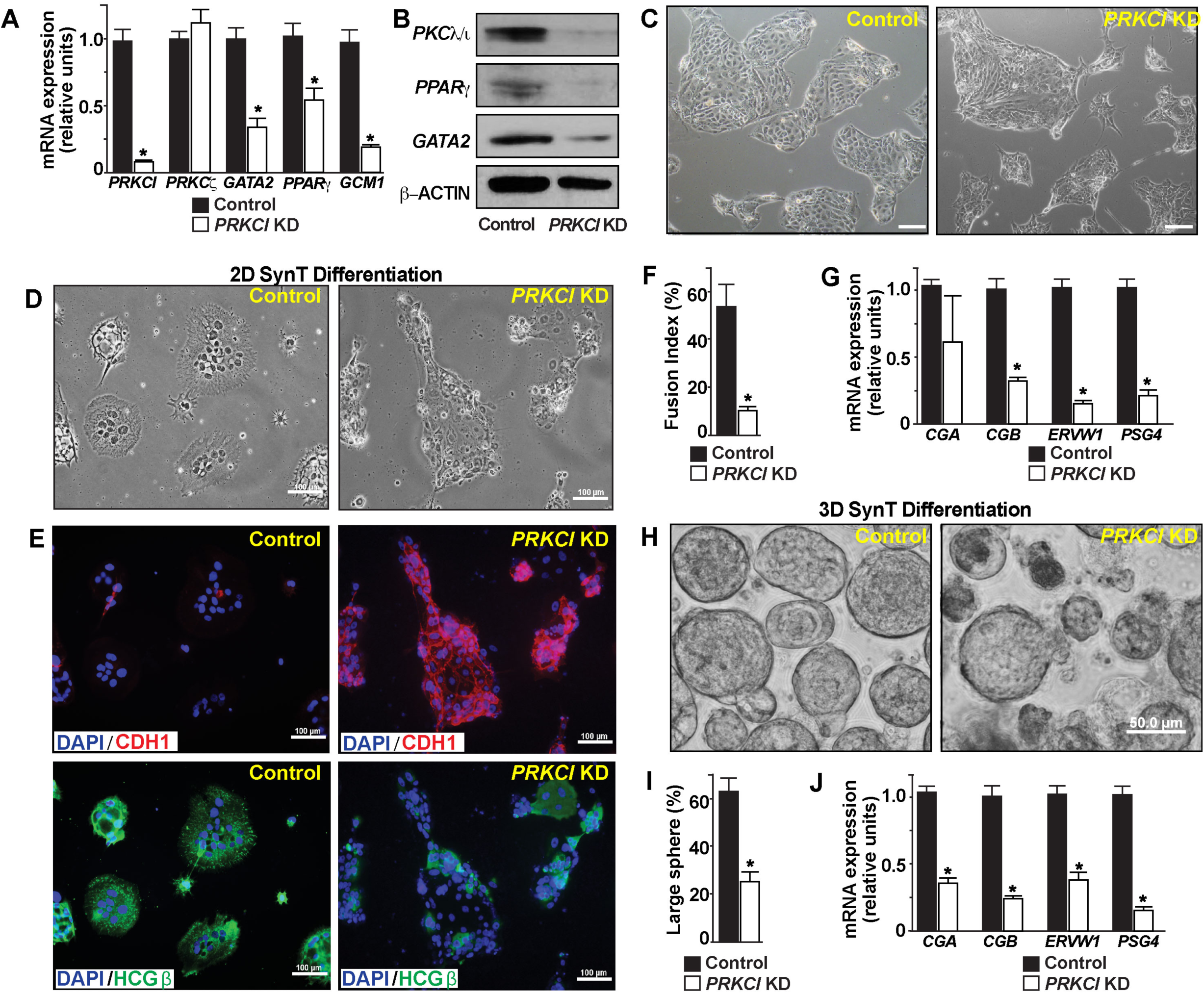
Loss of PKCλ/I impairs SynT differentiation potential in human TSCs. (A) Quantitative RT-PCR analyses showing loss of *GATA2, PPARG and GCM1* mRNA expression (mean ± SE; n = 4, p≤0.01) upon knock down of *PRKCI* in human TSCs (*PRKCI* KD human TSCs). The mRNA expression of *PRKCZ* was not affected. (B) Western blots show loss of PKCλ/I, GATA2 and PPARγ protein expressions in *PRKCI* KD human TSCs. (C) Micrographs showing morphology of control and *PRKCI* KD human TSCs. (D) and (E) Control and *PRKCI* KD human TSCs were subjected for 2-dimensional (2D) SynT differentiation on collagen coated adherent cell culture dishes. Image panels show altered cellular morphology (D); maintenance of E-Cadherin (CDH1) expression (Upper right panel in E); and impaired induction of HCGβ expression (lower right panel in E) in *PRKCI* KD human TSCs. (F) Cell fusion index was quantitated in control and *PRKCI* KD human TSCs. Fusion index was determined by measuring number of fused nuclei with respect to total number of nuclei within image fields (5 randomly selected fields from individual experiments were analyzed, 3 independent experiments were performed) (G) Quantitative RT-PCR analyses showing significant (mean ± SE; n = 4, p≤0.001) downregulation of mRNA expressions of SynT-specific markers in *PRKCI KD* human TSCs, undergoing 2D SynT differentiation. (H) Control and *PRKCI* KD human TSCs were subjected to 3-dimensional (3D) SynT differentiation on low attachment dishes. Micrographs show defective SynT differentiation, as assessed from formation of large cell-spheres, in *PRKCI* KD human TSCs. (I)- Quantification of 3D SynT differentiation efficiency was done by counting large cell-spheres (>50µm) from multiple fields (3 fields from each experiment, 3 individual experiments) (J) Quantitative RT-PCR analyses (mean ± SE; n = 4, p≤0.001) reveal impaired induction of SynT markers in *PRKCI KD* human TSCs, undergoing 3D SynT differentiation.

We cultured control and *PRKCI* KD human TSCs with forskolin on both high and low-adhesion culture plates to test the efficacy of both 2D and 3D syncytium formation. We assessed 2D SynT differentiation by monitoring elevated mRNA expressions of key SynT-associated genes, such as the HCGβ components *CGA, CGB;* retroviral fusogenic protein *ERVW1* and pregnancy-associated glycoprotein, *PSG4.* We also tested HCGβ protein expression and monitored cell syncytialization via loss of E-CADHERIN (CDH1) expression in fused cells. We found strong impairment of SynT differentiation of *PRKCI* KD human TSCs (Fig. 6D-6F). Unlike in control human TSCs, mRNA induction of key SynT-associated genes (Fig. 6G) as well as HCGβ protein expression were strongly inhibited in *PRKCI* KD human TSCs. Furthermore, *PRKCI* KD human TSCs maintained strong expression of CDH1 and showed a near complete inhibition of cell-fusion (Fig. 6E).

The impaired SynT differentiation potential in *PRKCI* KD human TSCs were also evident in the 3D culture conditions. Unlike control human TSCs, which efficiently formed large cyst-like spheres (larger than 50µm in diameter), *PRKCI* KD human TSCs failed to efficiently develop into larger spheres and mainly developed smaller cellular aggregates (Fig. 6H and 6I). Also, comparative mRNA expression analyses indicated more abundance of *PRKCI* mRNA in a few larger spheres, which were developed with *PRKCI* KD human TSCs, indicating that the large spheres are formed from cells, in which RNAi-mediated gene depletion was inefficient. We also confirmed inefficient induction of *CGA, CGB, ERVW1* and *PSG4* expressions in the small cell aggregates, which showed significant downregulation in *PRKCI* mRNA expressions (Fig. 6J), Thus, both the 2D and 3D SynT differentiation systems revealed impaired SynT differentiation potential of the *PRKCI* KD human TSCs.

### PKCλ/I signaling is essential for in vivo SynT differentiation potential of Human TSCs

As discussed above, our in-vitro differentiation analysis indicated an important role of PKCλ/I signaling in inducing SynT differentiation potential in the human TSCs. However, the *in vitro* differentiation system lacks the complex cellular environment and regulatory factors that control SynT development during placentation. Okae et al. showed that upon subcutaneous injection into non-diabetic-severe combined immunodeficiency mice (NOD-SCID mice), human TSCs invaded the dermal and underlying tissues to establish trophoblastic lesions (Okae et al., 2018). These trophoblastic lesions contain cells that represent all cell types of a villous human placenta, namely CTB, SynT and EVT. Therefore, we next tested *in vivo* SynT differentiation potential of *PRKCI* KD human TSCs via transplantation in the NOD-SCID mice (Fig. 7A). Upon transplantation, both control and *PRKCI* KD human TSCs generated tumors with similar efficiency (Fig. 7B) and the presence of human trophoblast cells were confirmed via analyses of human KRT7 expression (Fig. 7C). However, unlike control human TSCs, lesions that were developed from *PRKCI* KD human TSCs, were largely devoid of HCGβ-expressing SynT populations (Fig. 7D). Collectively, our *in vitro* differentiation and *in vivo* transplantation assays with human TSCs imply an essential and conserved molecular mechanism, in which the PKCλ/I signaling promotes expression of key transcription factors, like GATA2, PPARγ and GCM1 to assure CTB to SynT differentiation.

**Figure 7:**
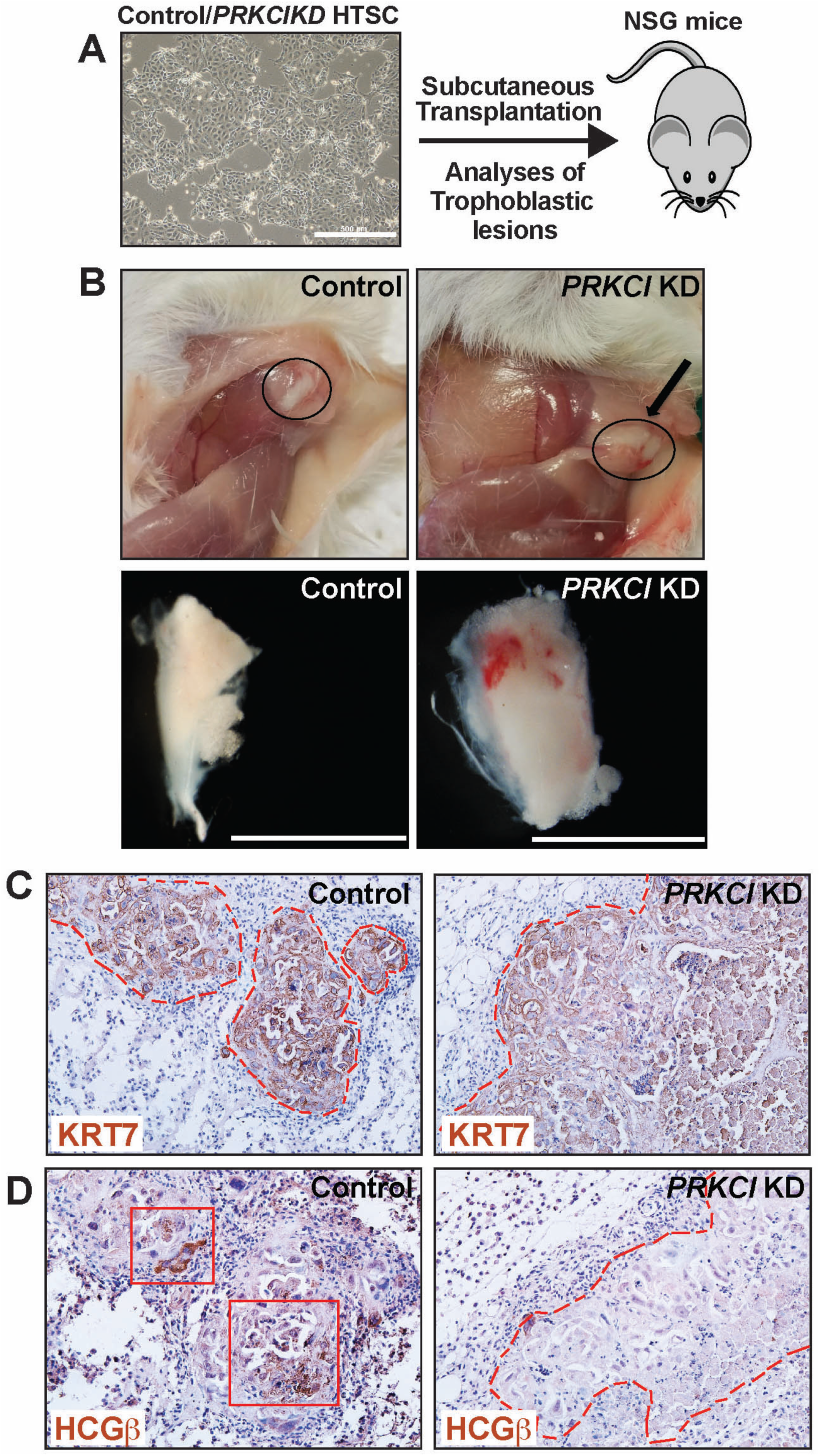
PKCλ/I signaling is essential for *in vivo* SynT differentiation potential of Human TSCs. (A) Schematics of testing *in vivo* developmental potential of human TSC via transplantation assay in NSG mice. Control and *PRKCI* KD human TSCs were mixed with matrigel and were subcutaneously injected into the flank of NSG mice. (B) Images show trophoblastic lesions that were developed from transplanted human TSCs. (C) and (D) Trophoblastic lesions were immunostained with anti-human cytokeratin7 (KRT7) antibody and anti-human HCGβ antibody, respectively. The trophoblastic lesions that were developed from both control and *PRKCI* KD human TSCs contained KRT7 expressing human trophoblast cells (C) but the lesions from *PRKCI* KD human TSCs lacked HCGβ expressing trophoblast populations (right panel in D), indicating impairment in SynT developmental potential.

## DISCUSSIONS

In this study, using both mouse knockout models and human TSCs we have uncovered an evolutionary conserved function of PKCλ/I signaling during trophoblast development and mammalian placentation. In recent years, placental development and the maternal-fetal interaction is being studied with considerable interest as defect in placentation in early post-implantation embryos lead to either pregnancy failure (Cockburn & Rossant, 2010; Pfeffer & Pearton, 2012; R. M. Roberts & S. J. Fisher, 2011; J. Rossant & J. C. Cross, 2001), or pregnancy-associated complications like IUGR and preeclampsia (Myatt, 2006; Pfeffer & Pearton, 2012; Redman & Sargent, 2005; J. Rossant & J. C. Cross, 2001), or serves as developmental causes for postnatal or adult diseases (Funai et al., 2005; Gluckman et al., 2008; Godfrey & Barker, 2000). Establishment of the intricate maternal-fetal relationship is instigated by development of the SynT populations, which not only establish the maternal-fetal exchange surface but also modulate the immune function and molecular signaling at the maternal-fetal interface to assure successful pregnancy (Aplin et al., 2017; Pavlicev et al., 2017). Our findings in this study indicate that PKCλ/I-regulated optimization of gene expression is fundamental to SynT development.

One of the interesting findings of this study is the specific requirement of PKCλ/I in establishing the SynT differentiation program. The lack of PKCλ/I expression in SynTs within first trimester and term human placentae indicates that PKCλ/I signaling is not required for maintenance of SynT function. The specific need of PKCλ/I to “prime” trophoblast progenitors for SynT differentiation is further evident from the phenotype of PKCλ/I-null mouse embryos. Although PKCλ/I is expressed in pre-implantation embryos and in TSPCs of placenta primordium, it is not essential for trophoblast cell lineage development at those early stages of embryonic development. Also, the development of TGCs in PKCλ/I-null placentae indicates that PKCλ/I is dispensable for the development of TGCs, a cell type that is equivalent to human EVTs. Rather, PKCλ/I is essential for labyrinth development at the onset of chorio-allantoic attachment, indicating a specific need of PKCλ/I in nascent SynT population.

Earlier studies with gene knockout mice indicated that both GCM1 and DLX3 are essential for the placental labyrinth development. The *Gcm1* knockout mice die at E10.5 (Schreiber et al., 2000) with major defect in SynT development. Similarly the defect in SynT development and placental labyrinth formation is evident in the *Dlx3* knockout mice by E9.5 (Morasso, Grinberg, Robinson, Sargent, & Mahon, 1999). Our findings in this study shows that the PKCλ/I signaling is essential to induce both *Gcm1 and Dlx3* transcriptions in the nascent SynT population. Not surprisingly, the PKCλ/I-KO embryos show a more severe phenotype with complete absence of SynTs in the developing placentae.

Our unbiased gene expression analyses in *Prkci* KD mouse TSCs strongly indicate that the PKCλ/I signaling ensures TSPC to SynT transition by maintaining expression of two conserved transcription factors, GATA2 and PPARγ. Both GATA2 and PPARγ are known regulators of SynT development. We showed that *Gcm1* and *Dlx3* are direct target genes of GATA2 in mouse TSCs (Pratik Home et al., 2017). Also loss of PPARγ in mouse TSCs is associated with complete loss of *Gcm1* induction during TSC to SynT differentiation (Parast et al., 2009). Thus, the impairment of *Gcm1 and Dlx3* induction upon loss of PKCλ/I-KO during TSPCs to SynT transition could be a direct result of the downregulation of GATA2 and PPARγ.

In this study, we also found that PKCλ/I-mediated regulation of GATA2, PPARγ and GCM1 is a conserved event in the mouse and human TSCs. In a recent study (Milano-Foster et al., 2019), we have shown that GATA2 regulates human SynT differentiation by directly regulating transcription of key SynT-associated genes, such as *CGA, CGB*, and *ERVW1*, via formation of a multi-protein complex, including histone demethylase KDM1A and RNA polymerase II, at their gene loci. Based on the conserved nature of GATA2 and PPARγ expression and their regulation by PKCλ/I in both mouse and human TSCs, we propose that a conserved PKCλ/I-GATA2/PPARγ-GCM1 regulatory axis instigates SynT differentiation during mammalian placentation. We also propose that the PKCλ/I-GATA2/PPARγ signaling axis mediates the progenitor to SynT differentiation by modulating global transcriptional program, which also involves epigenetic regulators, like KDM1A. As small molecules could modulate function of most of the members of this regulatory axis, it will be intriguing to test whether or not targeting this regulatory axis could be an option to attenuate SynT differentiation during mammalian placentation.

Another interesting finding of our study is impaired placental and embryonic development upon loss of PKCλ/I expression specifically in trophoblast cell lineage. The trophoblast-specific PKCλ/I depletion largely recapitulated the placental and embryonic phenotype of the global PKCλ/I-knockout mice. These results along with selective abundance of the PKCλ/I protein expression in TSPCs of post-implantation mouse embryos indicated that the defective embryo patterning in global PKCλ/I-knockout mice is probably an effect of impaired placentation in those embryos. It is well known that the cells of a gastrulating embryo has signaling cross talks with the TSPCs of a placenta primordium. Furthermore, defect in placentation is a common phenotype in many embryonic lethal mouse mutants. However, we have a poor understanding about how an early defect in labyrinth formation leads to impairment in embryo patterning and the specific phenotype of PKCλ/I-knockout mice provides an opportunity to better understand this process. Unfortunately, we do not have access to PKCλ/I-conditional knockout mouse model and the lentiviral-mediated shRNA delivery approach did not provide us with an option to deplete PKCλ/I in specific trophoblast cell types. During the completion of this study, we are also establishing a new mouse model, in which we will be able to conditionally delete PKCλ/I in specific trophoblast cell types to better understand how PKCλ/I functions in nascent SynT lineage and how it regulates the cross talk between the developing embryo proper and the developing placenta. Also, our finding in this study is the first known implication of PKCλ/I-signaling in human trophoblast lineage development and function. As defective SynT development could be associated with early pregnancy loss or pregnancy-associated disorders including fetal growth retardation, we also plan to study whether defective PKCλ/I function or associated downstream mechanisms are associated with early pregnancy loss and pregnancy associated disorders.

## EXPERIMENTAL PROCEDURES

### Animal and tissue collection

The *Prkci* knockout mice (B6.129-*Prkci*^*tm1Hed*^/Mmnc) were obtained from the Mutant Mouse Regional Resource Center, University of North Carolina (MMRRC UNC). The heterozygous animals were bred to obtain litters and harvest embryos at different gestational days. Pregnant female animals were identified by presence of vaginal plug (gestational day 0.5) and embryos were harvested at various gestational days. Uterine horns from pregnant females were dissected out and individual embryos were analyzed under microscope. Tissues for histological analysis were kept in dry-ice cooled heptane and stored at −80 for cryo-sectioning. Yolk sacs from each of the dissected embryos were collected and genomic DNA preparation was done using Extract-N-Amp tissue PCR kit (Sigma-XNAT2). Placenta tissues were collected in RLT buffer and RNA was extracted using RNAeasy Mini Kit (Qiagen – 74104). RNA was eluted and concentration was estimated using Nanodrop ND1000 spectrophotometer.

### Mouse TSC culture

Mouse TSCs were cultured using RPMI-1640 (Sigma) supplemented with 20% Fetal Bovine Serum (FBS), sodium pyruvate, β-mercaptoethanol and primocin (to avoid any mycoplasma contamination). Proliferative cells were maintained using mouse embryonic fibroblast (MEF)-conditioned media (CM) and basal media (70:30 ratio), FGF4 (25ng/ml) and heparin (1µg/ml). For differentiation assays, cells were allowed to differentiate by removing CM, FGF4 and heparin from the media. Cells were harvested at different time points and total RNA and protein were extracted for RT-PCR and western blot analyses.

### Human TSC culture

Human TSC lines, derived from first-trimester CTBs were described earlier (Okae et al., 2018). Although multiple lines were used, the data presented in this manuscript were generated using CT27 human TSC line. Human TSCs were cultured in DMEM/F12 supplemented with HEPES and L-glutamine along with cocktail of inhibitors. We followed the established protocol by Dr Arima’s group (Okae et al., 2018) and induced SynT differentiation using Forskolin. Both male and female human TSC lines were used for initial experimentation. To generate *PRKCI* KD human TSCs, shRNA-mediated RNAi were performed. For initial screening, both male and female cell lines were used. As we did not notice any phenotypic difference after PRKCI-depletion, a female *PRKCI* KD human TSC line and corresponding control set were used for subsequent experimentation.

### Human placental tissue sample analyses

Formaldehyde fixed, de-identified first trimester placental tissues were obtained from Mount-Sinai hospital, Toronto. Term Placental tissues were obtained at the University of Kansas Medical Center with consent from patients. All collections and Studies were approved by the University of Kansas IRB and the IRB at Mount Sinai Hospital. Fresh term placental tissues were embedded in OCT and cryo-sectioned. Formaldehyde fixed and frozen sections were analyzed by immunohistochemistry.

### mRNA expression analyses

Total RNA was extracted from the cells using RNeasy Mini Kit (Qiagen-74104) using manufacturer’s protocol. cDNA was prepared from total RNA (1000ng). Primer cocktail comprising of 200ng/µl oligo dT and 50ng/µl random hexamer was annealed to the RNA at 68° for 10 minutes, followed by incubation with the master mix comprising of 5X first strand buffer, 10mM dNTPs, 0.1M DTT, RNase Inhibitor and M-MLV transcriptase (200U/µl) at 42° for 1 hour. The cDNA solution was diluted to 10ng/µl and heat inactivated at 95 ° for 5 minutes. Real-time PCR was performed using oligonucleotides (listed in Table S4). 20ng equivalent of cDNA was used for amplification reaction using Power SYBR Green PCR master mix (Applied Biosystems-4367659). Primers are mentioned in supplemental information.

### Western blot analyses

Cell pellets were washed once with 1X PBS followed by addition of 1X Laemmli SDS-PAGE buffer for protein extract preparation and western blot analyses were performed following earlier described protocol (Saha et al., 2013). Antibodies are mentioned in supplemental information (Table S5).

### RNA interference with mouse and human TSCs

Lentiviral-mediated shRNA delivery approach was used for RNAi. For mouse cells, shRNA were designed to target the 3’ UTR sequence 5’-GTCGCTCTCGGTATCCTGTC-3’ of the mouse *Prkci* gene. For control, a scramble shRNA (Addgene-1864) targeting a random sequence 5’-CCTAAGGTTAAGTCGCCCTCGC-3’ was used. For human cells, short-hairpin RNA against *PRKCI* target sequence 5’-AGTACTGTTGGTTCGATTAAACTCGAGTTTAATCGAACCAACAGTACT-3’ was used to generate lentiviral particles (Sigma Mission-TRCN0000219727). Lentiviral particles were generated by transfecting HEK293T cells. Earlier described protocols (P. Home et al., 2009) were followed to collect and concentrate viral particles. Mouse and human TSCs were transduced using viral particles with equal MOI at 60-70% confluency. The cells were treated with 8µg/ml polybrene prior to transduction. Cells were selected in the presence of puromycin (1.5-2µg/ml). Selected cells were tested for knockdown efficiency and used for further analyses. Freshly knocked-down cells were used for each individual experimental set to avoid any silencing of shRNA expression due to DNA-methylation at LTR. To generate data at least four individual experiments were done to get statistically significant results.

### Trophectoderm-specific *Prkci* knockdown

Morula from day 2.5 plugged CD-1 superovulated females were treated with Acidic Tyrode’s solution for removal of zona pellucida. The embryos were immediately transferred to EmbryoMax Advanced KSOM media. Embryos were treated with viral particles having either control pLKO.3G empty vector or shRNA against *Prkci* for 5hours. The embryos were washed 2-3 times, subsequently incubated overnight in EmbryoMax Advanced KSOM media and transferred into day 0.5 pseudopregnant females the following day. Uterine horns of control and knockdown sets were harvested at day 9.5. Placental tissue was obtained for RNA preparation to validate knockdown efficiency. Embryos were either dissected or kept frozen for sectioning purpose.

### Immunofluorescence and Immunohistochemistry analyses

Immunofluorescence was performed using 10µm embryo cryosections. The sections were fixed using 4% paraformaldehyde in 1X PBS, permeabilized using 0.25% Triton X-100 in 1X PBS and blocked for 1 hour using 10% Normal Goat serum (Thermo Fisher scientific-50062Z). The details of antibodies used are listed in Table S5. Immunohistochemistry was performed using paraffin sections of human placenta. The slides were deparaffinized by histoclear and subsequently with 100%, 90%, 80% and 70% ethanol. Antigen retrieval was done using Decloaking chamber at 80°C for 15minutes. The slides were washed with 1X PBS and treated with 3% H_2_O_2_ to remove endogenous peroxidase followed by 3 times wash with 1X PBS. 10% goat serum was used as a blocking reagent for 1hour at RT followed by overnight incubation with 1:100 dilution of primary antibody or IgG at 4°C. The slides were washed with 1X PBS and 1:200 dilution of secondary antibody was used for 1 hour at RT. The slides were washed again with 1X PBS followed by treatment with horseradish peroxidase for 20 minutes at RT. The slides were washed again and proceeded to color development using DACO 1ml buffer and 1 drop of chromogen. The reaction was stopped in distilled water after sufficient color developed. The slides were counterstained with Mayer’s hematoxylin for 5minutes and washed with warm tap water until sufficient bluish coloration observed. The slides were then dehydrated by sequential treatment using 70%, 80%, 90%, 100% ethanol and histoclear. The sections were completely dried and mounted using Toluene as mountant and imaged using Nikon TE2000 microscope.

### Cell proliferation assay

Mouse and Human TSCs were seeded (30,000cells/well of 12 well plate) and cultured for 24, 48, 72, 96 hours to assess cell proliferation. Cell proliferation was assessed using both BrDU labeling assay and detection kit (Roche Ref#11296736001) in live cells and with MTT assay kit (Catalog# Sigma CGD1) after harvesting cells at distinct time intervals. We followed manufacturers’ protocol.

### RNA-In Situ hybridization (RNA-ISH)

RNA-ISH was performed using ACDBio RNASCOPE kit (Catalog # 320851 and 322310) and following manufacturer’s protocol. The probes for *Prkci* (Catalog# 403191) and *Dlx3* (Catalog# 425191) were custom designed by ACDbio.

### RNA-seq analyses

Total RNA was used to construct RNA-seq libraries using the Illumina TruSeq Stranded Total RNA Sample Preparation Kit according to manufacturer’s instructions. RNA seq was performed using Illumina HiSeq 2500 platform. Raw sequence reads in fastq format were mapped to the *Mus musculus* GRCm38 genome with STAR-2.5.2b (Dobin et al., 2013). Gene expression levels were estimated with StringTie-1.3.0 (Pertea et al., 2015) and cuffmerge (Trapnell et al., 2012) from the cufflinks2 (Anders, Pyl, & Huber, 2015) package. Gene counts were obtained with htseq-0.6.1p1 followed by differential gene expression analysis with EdgeR (Robinson, McCarthy, & Smyth, 2010).

### Transplantation of human TSCs into NSG mice

For transplantation analyses in NSG mice, Control and *PRKCI* KD human TSCs were mixed with matrigel and used to inject subcutaneously into the flank of NSG mice (6-9-week-old). 10^7^cells were used for each transplantation experiment. Mice were euthanized after 10 days and trophoblastic lesions generated were isolated, photographed, measured for size, fixed, embedded in paraffin and sectioned for further analyses.

## ACKNOWLEDGEMENTS

This research was supported by NIH grants HD062546, HD0098880, HD079363, a bridging grant support under the Kansas Idea Network Of Biomedical Research Excellence (K-IBRE, P20GM103418) to Soumen Paul, a University of Kansas Biomedical Research Training Program grant to Bhaswati Bhattacharya and a NIH Center of Biomedical Research Program pilot grant to Pratik Home. This study was supported by various core facilities, including the Genomics Core, Transgenic and Gene Targeting Institutional Facility, Imaging and histology core facility and the Bioinformatics core of the University of Kansas Medical Center.

## CONTRIBUTIONS

Soumen Paul and Bhaswati Bhattacharya conceived and designed the experiments and wrote manuscripts: Bhaswati Bhattacharya, Pratik Home, Avishek Ganguly, Ananya Ghosh, Md. Rashedul Islam and Soma Ray performed experiments. Sumedha Gunewardena performed bioinformatics analyses. Courtney Marsh, Valerie French, Hiroaki Okae and Takahiro Arima provided reagents.

