## Supplementary Information for "Atypical Protein Kinase C iota (PKCλ/ι) Ensures Mammalian Development by Establishing the Maternal-Fetal Exchange Interface"

The supplementary information contains the following items.

(i) Supplementary Figures
(ii) Legends for Supplementary Figures
(iii) Supplementary Tables containing oligomers and antibodies

(Additional supplementary tables, containing gene expression data from RNA-seq analyses were uploaded as Excel file)

### Supplementary Figure1

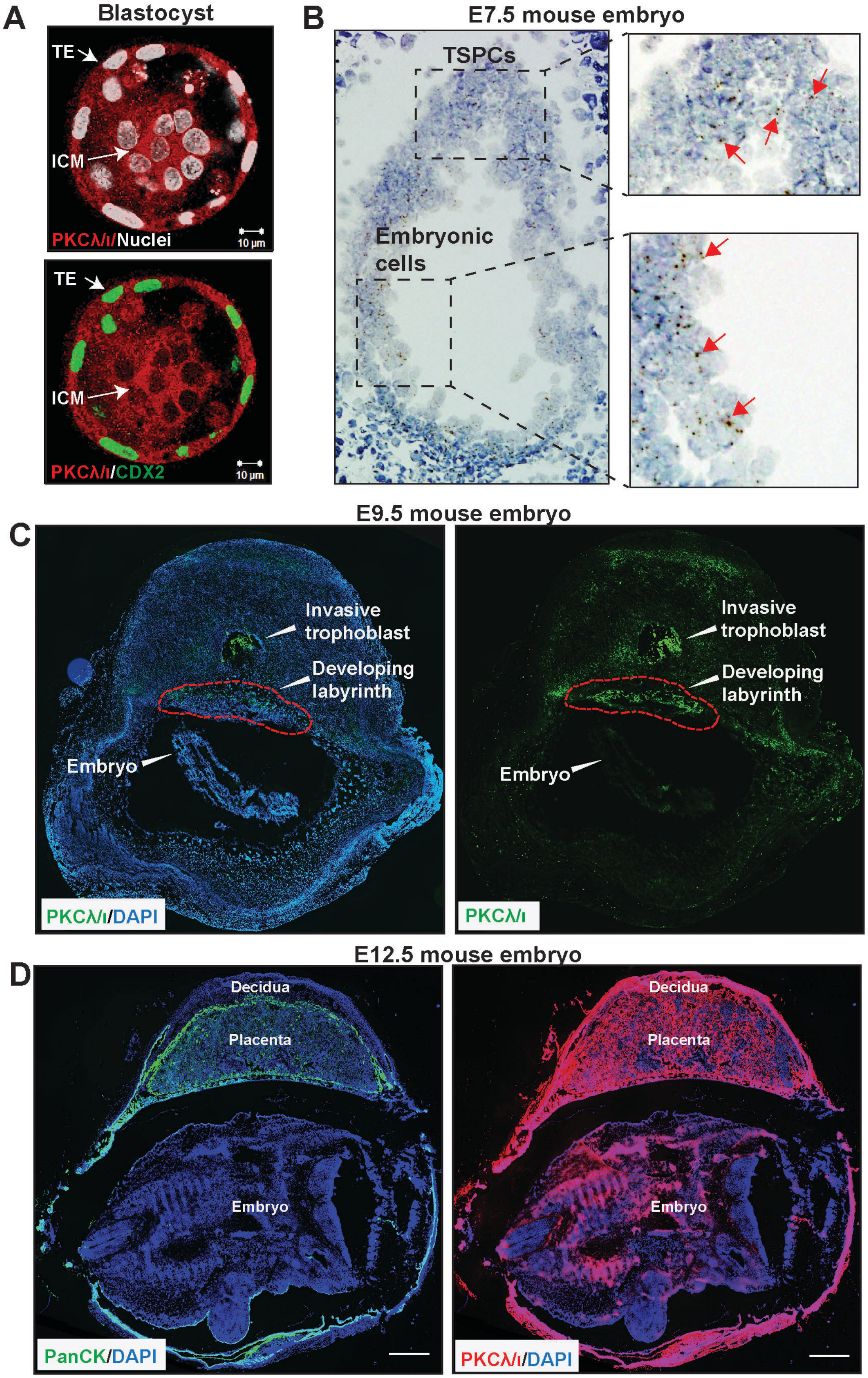

### Supplementary Figure 2

**A**

E7.5 dissection

| Total no of embryos analyzed: 86 (12 litters) | | |
|---|---|---|
| No of embryos | | Average length of embryo |
| WT | 16 | 1±0.1mm |
| HET | 50 | 1±0.1mm |
| KO | 20 | 1±0.1mm |

E9.5 dissection

| Total no of embryos analyzed: 66 (8 litters) | | |
|---|---|---|
| No of embryos | | Embryos having smaller placenta |
| WT | 14 | 0 |
| HET | 37 | 1 |
| KO | 15 | 15 |

**B** Gene expression analyses in E9.5 placentae

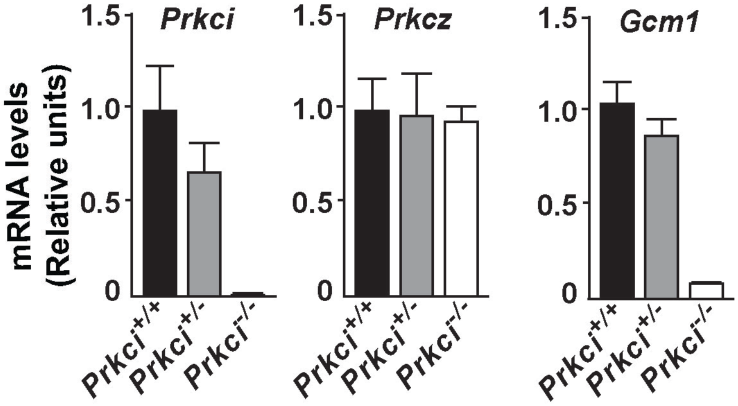

#### Supplementary Figure 3

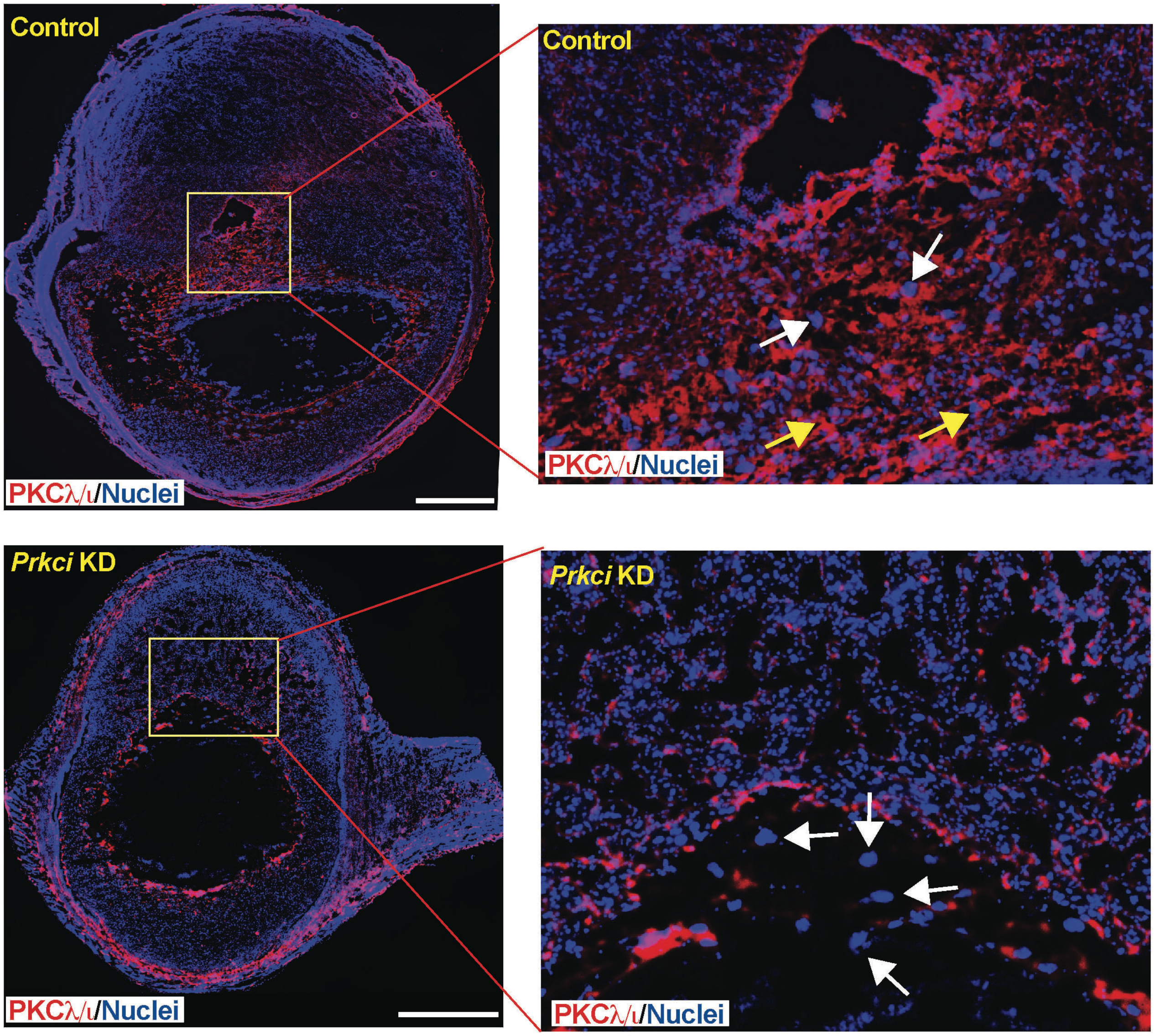

#### Supplementary Figure 4

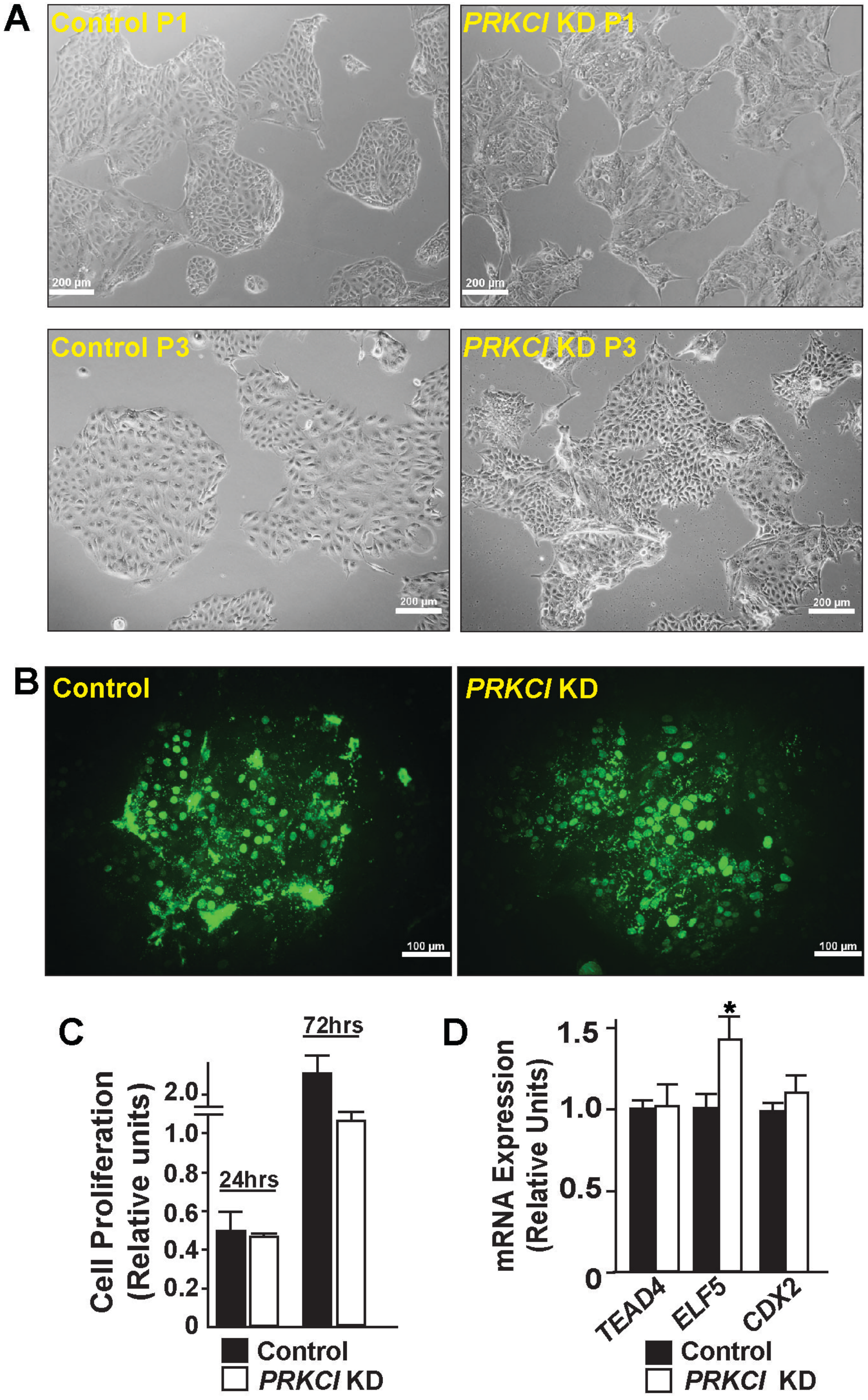

#### SUPPLEMENTARY FIGURE LEGENDS

**Supplementary figure 1.** (A) Immunofluorescence of mouse blastocyst using anti-PKCλ/ι antibody (red), anti-CDX2 (green) and DAPI. Images show that PKCλ/ι is expressed in both inner cell mass and in trophectoderm. (B) RNA in situ hybridization was performed using fluorescent probes against *Prkci* mRNA in E7.5 mouse implantation site. Images show expression of *Prkc*i mRNA (red punctate dots marked by red arrows) in the embryonic cells as well in the TSPCs within placenta primordium. (C) RNA in situ hybridization was performed in E9.5 mouse implantation site showing prominent *Prkc*i mRNA expression in the labyrinth zone and maternal decidua. (D) Immunofluorescence of E12.5 mouse embryo using anti-pan-cytokeratin (PanCK, trophoblast marker) antibody (left panel) and anti-PKCλ/ι antibody (right panel). Nuclei were stained with DAPI. At this developmental stage, PKCλ/ι protein expression is prominent in both placenta and in the embryo proper.

**Supplementary figure 2.** (A) Tables showing total number of embryos analyzed at E7.5 and E9.5 stages. (B) Quantitative RT-PCR analyses using placental tissue from control and PKCλ/ι KO embryos. *Prkcz* mRNA level remained unaltered in KO placentae. PKCλ/ι KO placentae showed near complete loss of *Gcm1*, a SynT-specific gene.

**Supplementary figure 3.** Control and *Prkci* KD mouse embryos were immunostained with anti-PKCλ/ι antibody (red) and DAPI. PKCλ/ι expression was significantly reduced in the TGCs of *Prkci* KD placenta (marked by white arrows).

In contrast high level of PKCλ/ι expression was maintained in trophoblast cells of control placenta (marked by yellow arrows).

**Supplementary figure 4.** (A) Micrographs show maintenance of stem state colony morphology of both control and *PRKCI* KD human TSCs for multiple passages. (B) Immunofluorescence images show incorporation of BrdU in both control and *PRKCI* KD human TSCs, indicating maintenance of cell proliferation (C) Quantitative Assessment of cell proliferation rate of control and *PRKCI* KD human TSCs by MTT assay. (D) Quantitative RT-PCR analyses (mean ± SE; n = 4, p≤0.01) showing relative mRNA expression of trophoblast stem state genes in control and *PRKCI* KD human TSCS.

#### Supplementary Tables for Oligomers and Antibodies

##### *Table S4: Oligomers Used for The Study*

###### *Genotyping primers:*

| Gene | Forward | Reverse |
|---|---|---|
| *Prkci wildtype* | *AACACCAGGGAGAGTGG* | *GCCTAGAAAGTAACCCAC* |
| *Prkci knockout* | *AACACCAGGGAGAGTGG* | *GACGAGTTCTTCTGAGGGG* |

###### *RT-PCR mouse primers*

| Gene | Forward | Reverse |
|---|---|---|
| *Prkci Ex9* | *CTAGGTCTGCAGGATTTCG* | *TTGACGAGCTCTTTCTTCAC* |
| *Prkci (Ex1/2)* | *TCCGGGTGAAAGCCTACTAC* | *CAAAGGAGATGGAAGGCTCA* |
| *Prkcz* | *GGACAACCCTGACATGAACAC* | *GGCCTTGACAGACAGGAAAC* |
| *18SrRNA* | *AGTTCCAGCACATTTTGCGAG* | *TCATCCTCCGTGAGTTCTCCA* |
| *Cdx2* | *GGACGTGAGCATGTATCCTAGCT* | *TAACCACCGTAGTCCGGGTACT* |
| *Eomes* | *ACCAATAACAAAGGTGCAAACAAC* | *TGGTATTTGTGCAGAGACTGCAA* |
| *Esrrb* | *AGTACAAGCGACGGCTGG* | *CCTAGTAGATTCGAGACGATCTTAGTCA* |
| *Elf5* | *ATGTTGGACTCCGTAACCCAT* | *GCAGGGTAGTAGTCTTCATTGCT* |
| *Gata3* | *CGGGTTCGGATGTAAGTCGA* | *GTAGAGGTTGCCCCGCAGT* |
| *Gcm1* | *AGAGATACTGAGCTGGGACATT* | *CTGTCGTCCGAGCTGTAGATG* |
| *Dlx3* | *CACTGACCTGGGCTATTACAGC* | *GAGATTGAACTGGTGGTGGTAG* |
| *Tead4* | *ATCCTGACGGAGGAAGGCA* | *GCTTGATATGGCGTGCGAT* |
| *Gata2* | *GGAAGATGTCCAGCAAATCC* | *TGGAGAGCTCCTCGAAACAT* |
| *Pparg* | *AGCTGTCATTATTCTCAGTGGAG* | *ATGTCCTCGATGGGCTTCAC* |
| *Ascl2* | *AAGCACACCTTGACTGGTACG* | *AAGTGGACGTTTGCACCTTCA* |
| *Hand1* | *CTACCAGTTACATCGCCTACTTG* | *ACCACCATCCGTCTTTTTGAG* |
| *Cx31* | *TCTGGCTGTCAGTAGTGTTCG* | *GCCTGGTGTTACAGTCAAAGTC* |
| *Tpbpa* | *TCCGGTCAGCTAACTGATGA* | *TCCTCTTCAAACATTGGGTGT* |
| *Prl3d1* | *ACATTTATCTTGGCCGCAGATGTGT* | *TTTAGTTTCGTGGACTTCCTCTCGAT* |

###### *RT-PCR human primers:*

| Gene | Forward | Reverse |
|---|---|---|
| *PRKCI* | *AGGTCCGGGTGAAAGCCTA* | *TGAAGAGCTGTTCGTTGTCAAA* |
| *PRKCZ* | *ATGACGAGGATATTGACTGGGT* | *CAGGAGTGTAATCCGACCAGG* |
| *GATA2* | *CCAGCTTCACCCCTAAGCAG* | *CCACAGTTGACACACTCCCG* |
| *PPARG* | *ACCAAAGTGCAATCAAAGTGGA* | *ATGAGGGAGTTGGAAGGCTCT* |
| *GCM1* | *GGCGCAAGATCTACCTGAGA* | *CACAGTTGGGACAGCGTTT* |
| *ERVW-1* | *CTACCCCAACTGCGGTTAAA* | *GGTTCCTTTGGCAGTATCCA* |
| *CGA* | *TCTGGTCACATTGTCGGTGT* | *TTCCTGTAGCGTGCATTCTG* |
| *CGB* | *GTGTGCATCACCGTCAACAC* | *GGTAGTTGCACACCACCTGA* |
| *PSG4* | *CGATGGGACTGGAGGAGTAA* | *AGTTGCTGCTGGAGATGGAG* |
| *HPRT1* | *ACCCTTTCCAAATCCTCAGC* | *GTTATGGCGACCCGCAG* |
| *18SrRNA* | *AACCCGTTGAACCCCATT* | *CCATCCAATCGGTAGTAGCG* |
| *CDX2* | *CTGGTTTCAGAACCGCAGAG* | *TGCTGCTGCAACTTCTTCTT* |
| *ELF5* | *CTGCCTTTGAGCATCAGACA* | *TCCAGTATTCAGGGTGGACTG* |
| *TEAD4* | *ACGGCCTTCCACAGTAGCAT* | *CTTGCCAAAACCCTGAGACT* |

***Table S5: Antibodies Used for the Study***

***List of primary antibodies:***

| **Name** | **Company** | **Catalog#** |
|---|---|---|
| *PKCiota* | *BD Transduction Laboratories* | *610175* |
| *Pan-cytokeratin* | *Abcam* | *ab9377* |
| *CDX2* | *Abcam* | *ab76541* |
| *β-actin* | *Sigma* | *A5441* |
| *HCGβ* | *Abcam* | *ab53087* |
| *Cytokeratin 7* | *Dako* | *M7018* |
| *E-cadherin* | *Abcam* | *ab1416* |
| *PPARγ (mouse)* | *Proteintech* | *16643-1-AP* |
| *PPARγ (human)* | *SantaCruz* | *sc81152* |
| *GATA2* | *Abcam* | *ab109241* |

***List of secondary antibodies:***

| **Name** | **Company** | **Catalog#** |
|---|---|---|
| *Alexa fluor 488 goat anti-rabbit IgG* | *Invitrogen* | *A11008* |
| *Alexa fluor 568 goat anti-mouse IgG* | *Invitrogen* | *A11031* |
| *Alexa fluor 488 donkey anti-mouse IgG* | *Invitrogen* | *A21202* |
| *Alexa fluor 568 donkey anti-rabbit IgG* | *Invitrogen* | *A10042* |
| *Goat anti-mouse IgG-HRP* | *Santa Cruz* | *sc2005* |
| *Goat anti-rabbit IgG-HRP* | *Santa Cruz* | *sc2004* |
